# Multiplexed whole animal imaging with reversibly switchable optoacoustic proteins

**DOI:** 10.1101/2020.02.01.930222

**Authors:** Kanuj Mishra, Mariia Stankevych, Juan Pablo Fuenzalida-Werner, Simon Grassmann, Vipul Gujrati, Uwe Klemm, Veit R. Buchholz, Vasilis Ntziachristos, Andre C. Stiel

## Abstract

We describe two photochromic proteins for cell-specific *in vivo* optoacoustic (OA) imaging with signal unmixing in the temporal domain. We show highly sensitive, multiplexed visualization of T lymphocytes, bacteria and tumors in the mouse body and brain. We developed machine learning-based software that allows commercial imaging systems to be used for temporal unmixed OA imaging, enabling its routine use in life sciences.

## Introduction

Photo- or optoacoustic (OA) imaging combines optical contrast with ultrasound resolution, enabling high-resolution, real-time *in vivo* imaging well beyond the 1 mm penetration depth typical of microscopy methods^1,2^. OA already has provided intriguing insights into tumor heterogeneity^3^, Crohn’s disease^4^, neuronal dynamics^5^, psoriasis^6^ and brown fat metabolism^7^ based on endogenous contrast from hemoglobin and lipids^8,9^. However, OA imaging has not yet become a routine tool in life sciences because of the lack of strong OA contrast agents that can be expressed in desired cell types^10^. The few transgenic labels used in OA so far^9^ give weak signals that cannot rise above the strong background due to hemoglobin. Photochromic proteins that can be reversibly switched between two states by light can overcome this limitation by entirely separating the label signal, which modulates in accordance with the illumination, from the background, which remains constant (Fig. 1A)^11^. This concept, despite being validated in several studies^12–17^, has not been implemented widely because it requires complex instrumentation and data analysis tools. Here we introduce two reversibly switchable OA proteins (rsOAPs) and demonstrate their use with widely accessible off-the-shelf commercial imaging systems as well as our open-access machine-learning (ML) based software code for analysis. One of our new rsOAPs shows high switching speeds and dynamic range of photo-modulation that allow us to resolve the signals of different rsOAPs in close proximity in the same animal, demonstrating the potential for simultaneous tracking of different cellular processes through temporal multiplexing.

**Fig. 1:**
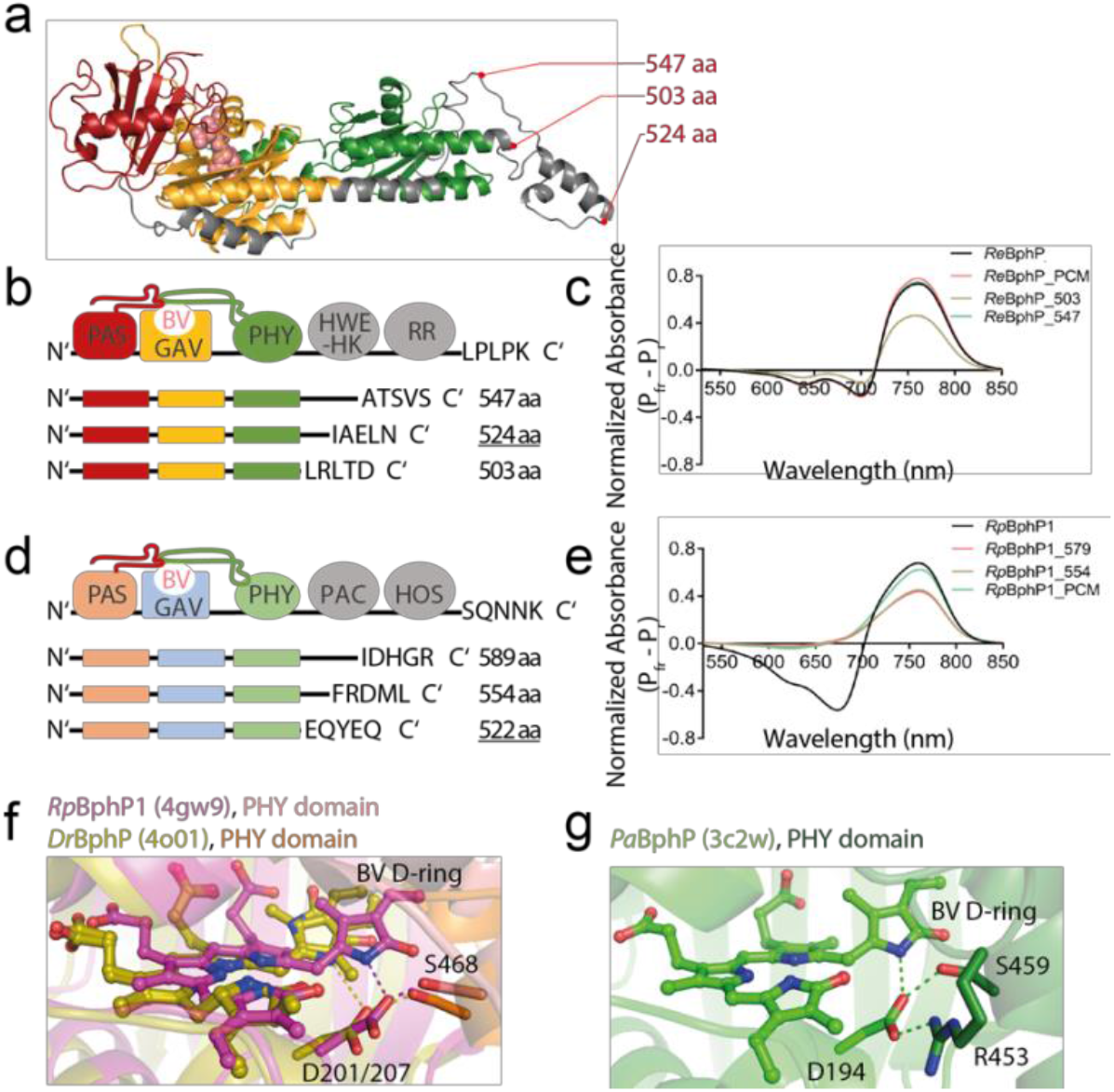
Structure guided design of rsOAPs. **a** Homology model (iTasser, based on 6g1y) of *Re*BphP-PCM. Truncation sides indicated. **b** Schematic representation of truncations. **c** Photo-induced differential spectra for truncations. **d** and **e** Similar representations for *Rp*BphP1. **f** Stabilization of the BV D-ring in *Rp*BphP1 and *Dr*BphP. **g** Similar representation for PaBphP which shows an arginine similar to *Re*BphP presumably abstracting D194 and destabilizing the P_fr_ state yielding a faster photo-switching.

## Results

### Engineering of rsOAPs

Bacterial photoreceptors called bacteriophytochromes (BphPs) 18 have emerged as most suitable for rsOAP development due to their strong absorption in the near-infrared range and low photo-fatigue^19^. To identify the most promising candidate for further development, we screened eight native BphPs (Suppl. Table 1) and selected the one from *Rhizobium etli*. A set of truncations enabled us to minimize its size and optimize its photo-switching characteristics. In brief, based on existing structural data as well as homology models we created truncations containing the minimum PAS-GAF-PHY photosensory core domains (PCM = photosensory core module) together with extra amino acids from the annotated linkers between PHY and Histidine Kinase domains and tested their characteristics in regard to signal generation and photo-switching (Fig 1a-e, Suppl. Note 1). The final variant *Re*BphP-PCM shows 2-fold larger change in OA signal (Fig. 2g), >5-fold faster switching (Fig. 2c and d) and greater resistance to photo-fatigue than other rsOAPs (Fig. 2e) while its high molar absorbance is on par with the recently described *Deinococcus radiodurans Dr*BphP-PCM^16^ (92,000 M^-1^ cm^-1^, Fig. 2b and 2g). Those characteristics enable higher numbers of switching cycles per second, which improves sensitivity and allows imaging over longer timeframes. On the molecular level this acceleration of switching speed is the result of a less stabilized P_fr_ state favouring the photoinduced transition to P_r_. The destabilization is likely caused by an arginine present in *Re*BphP but not in *Rp*BphP1 and *Dr*BphP. This arginine, by interacting with a conserved aspartate which in turn interacts with the D-ring of the P_fr_ state chromophore, weakens P_fr_ stabilization. (Fig 1f and g, Suppl. Note 1).

**Fig. 2:**
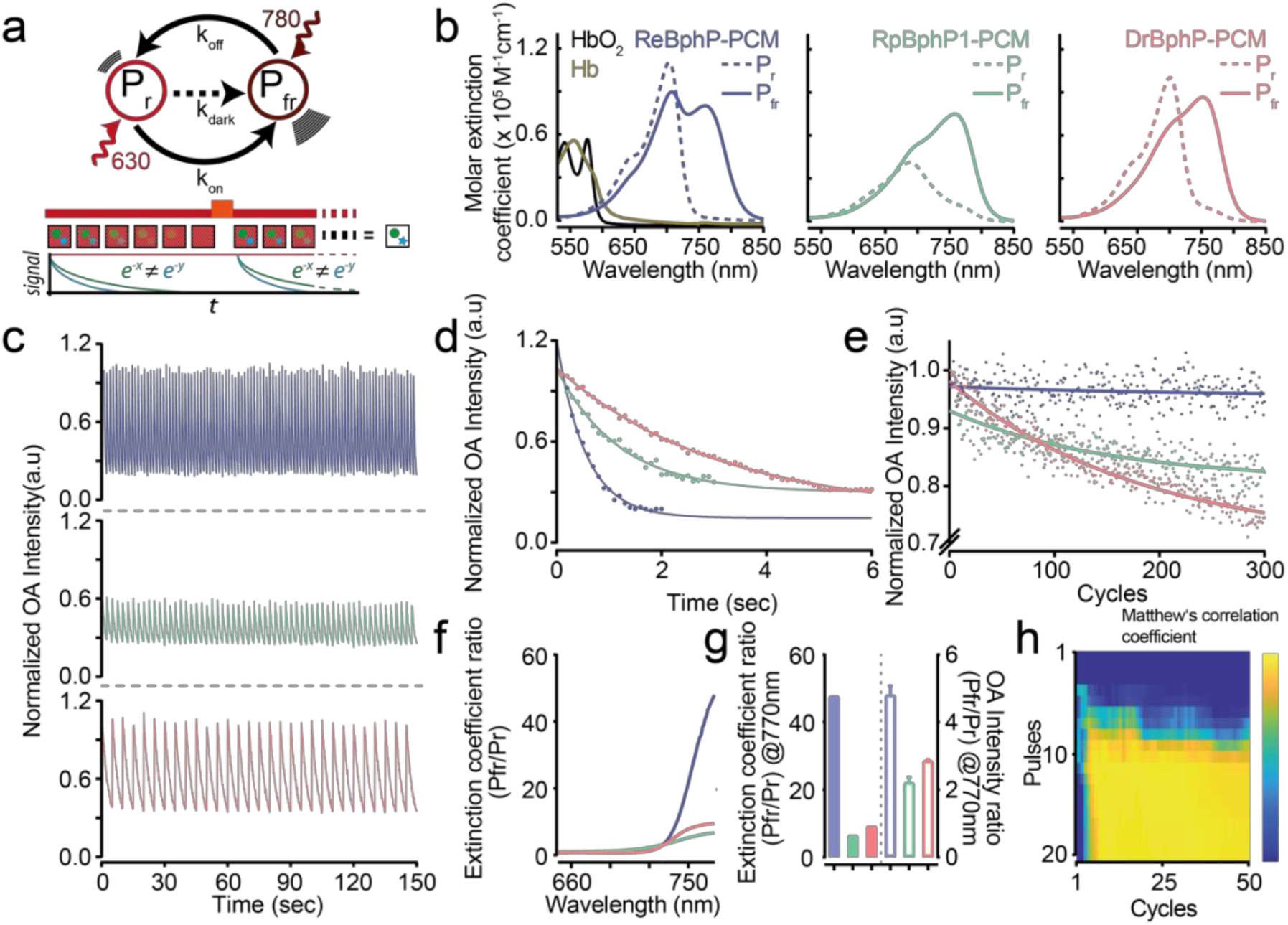
*In vitro* characterization of *Re*BphP-PCM and *Rp*BphP1-PCM in comparison to another rsOAP *Dr*BphP-PCM. **a** Principle of photo-switching in BphP-derived rsOAPs (top) and concept of temporal unmixing of two labels (green ball and blue star; bottom. The illumination is shown as dark-red (780 nm) and red (630 nm) bar. P_r_ refers to the red state, while P_fr_ refers to the far-red state. The bottom part of the panel was adapted with permission from^11^, Copyright 2019 American Chemical Society. **b** Absorbance spectra of P_r_ and P_fr_ states of the three rsOAPs used in this study in comparison to oxy- and deoxygenated hemoglobin (HbO_2_ and Hb). Spectra for hemoglobin were taken from 1999, Scott Prahl, omlc.org. **c** Switching cycles of the rsOAPs. Only the OA signal at 770 nm is shown. The number of pulses per cycle was chosen based on the time constants of the proteins. **d** Single switching-cycle from panel c, shown with an exponential fit. The dependence of switching kinetics on light fluence and laser repetition rate can be found in Suppl. Fig. 6. **e** Photo fatigue of the proteins per cycle. The number of pulses per cycle was chosen based on the time constants of the proteins. **f** Absorbance ratio between the P_fr_ and P_r_ state for different wavelengths. **g** Absorbance (filled bars) and OA signal intensity (hollow) ratio between the P_fr_ and P_r_ state for the three rsOAPs at 770 nm. **h** Matthew’s coefficient shown as a function of number of cycles and pulses. Shown is the analysis of a 4T1 tumor expressing *Re*BphP-PCM (see below); histology was used as ground truth (see Supp. Figure 6 and Suppl. Note 2 for details). All proteins have been adjusted to equal Soret peak absorption.

Our truncation strategy also proved successful in obtaining a switchable *Rp*BpliPI-PCM from *Rhodopseudomonas palustris*, in contrast to a previous report that truncated forms of this protein do not undergo reversible switching ^16^. Our engineered *Rp*BphP1-PCM maintains the far-red state (P_fr_) extinction coefficient and photochromic behaviour of the parental *Rp*BphP1 (Fig. 2b and 1g, Suppl. Fig. 1), and the change in its OA signal following illumination at 770 nm is similar to that of the previously described *Dr*BphP-PCM (Fig. 2g). (Plasmids for expressing rsOAPs in bacteria, eukaryotic cells or for introduction into viral vectors can be obtained from Addgene.) Both new rsOAPs are monomeric (Suppl. Fig. 2) and show higher expression in mammalian cells than the full-length parental proteins (Suppl. Fig. 3). The two developed rsOAPs and *Dr*BphP-PCM show distinctive switching speeds which is the reason for our ability to discriminate the proteins *in vivo* successfully. As a result, probes expressed in different cells in close proximity in the animal can be distinguished during high-resolution OA imaging.

### OA imaging using ML based temporal unmixing

We performed all OA imaging using an off-the-shelf, commercially available multispectral OA tomography device (Fig. 4a) with a 10-Hz pulsed tunable laser and a 256-element transducer array (MSOT, iThera Medical). Off-switching of rsOAPs was achieved with light at 770 nm, which gave the highest difference in OA signal intensity between the “on” and “off” states (Suppl. Fig. 5a), while on-switching was achieved using light at 680 nm. Lower wavelengths did not substantially improve the transition to the “on” state (Suppl. Fig. 5b). The number of laser pulses per wavelength was chosen to cover the full switching kinetics, but it can be significantly reduced for time-resolved studies using information-content analysis which allows an estimate of the minimal number of cycles and pulses per cycle required to discern the labeled structure, thus, effectively limiting imaging dwell time, which is essential for e.g. time-resolved studies (Fig. 2h, Suppl. Fig. 6, Suppl. Note 2). All temporal unmixing was conducted with in-house code developed to analyze time-varying patterns in the reconstructed data in the frequency and time domains using classic machine learning approaches (Fig. 3, Methods and Suppl. Note 3 and 4). In brief, after running fluence and motion correction on the data a range of distinctive features was extracted from the photo-modulated signal for each voxel of the tomography images. Based on a set of such data and corresponding histology as ground truth a bagged random forest algorithm^20^ was trained and validated on independent datasets of a different type to prevent over-fitting. The ensuing model was then used to analyze all data in this study. The code for data preparation, for analysis with the model used in this work and for generation of new models is available to the community along with graphical user interfaces.

**Fig. 3:**
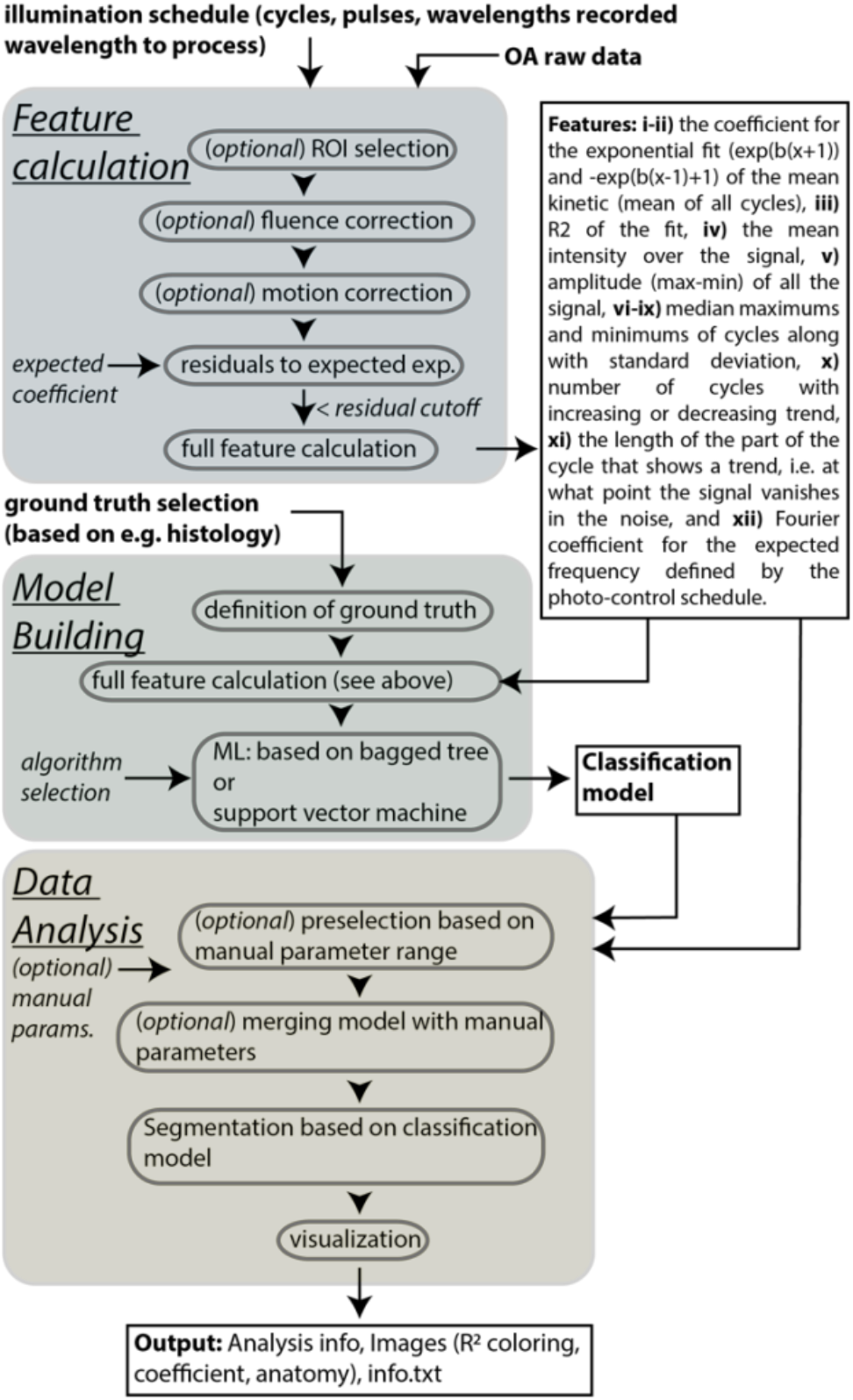
Schematic flow of the ML based analysis strategy and main scripts. The time varying patterns in the OA raw data are extracted in the “Feature calculation” (blue) and analyzed using a classification model in the “Data Analysis” step (yellow). In “Model building” (green) a classification model is trained based on imaging data with associated histology ground truth. In the script two algorithms can be selected: bagged tree or support vector machine. For uniformity the images shown in this work exclusively use the bagged tree approach although the support vector machine has some virtues (Suppl. Note 4).

First, we used rsOAPs for superficial *in vivo* imaging. We imaged the development of 4T1 mouse mammary gland tumors co-expressing *Re*BphP-PCM and GFP after they were grafted onto the backs of FoxN1 nude mice (n=3). The initial population of 0.8 x 10^6^ injected cells was readily visualized separate from all background absorbers (Suppl. Fig. 7a-d) as was the growing tumor mass at all days after injection (Fig. 4b). To test whether such imaging is also possible in brain tissue after light passes through the skull, we implanted 0.7 x 10^6^ 4T1 cells coexpressing *Re*BphP-PCM and GFP at a depth of 3.6 mm and imaged them immediately thereafter. Comparison of the OA images with fluorescence images obtained after sacrificing mice revealed perfect overlap of the labelling, confirming background-free identification of 1.4 x 10^5^ cells/ deep in the mouse brain (Fig. 4c). Next, we used the same rsOAP to image deep-seated tumors of HCT116 human colon carcinoma cells implanted intraperitoneally (n=2). From day 7 onward we were able to visualize the growth of several individual tumor sites to a depth of ~1 cm (Suppl. Fig. 7e and f). Comparison of OA images and histology obtained after sacrifice confirmed identification of all malignant tissue (Fig. 4d-f and Suppl. Figure 7e and g), including small tumors or metastatic patches containing less than 10,000 cells (Suppl. Fig. 7i and j).

**Fig. 4:**
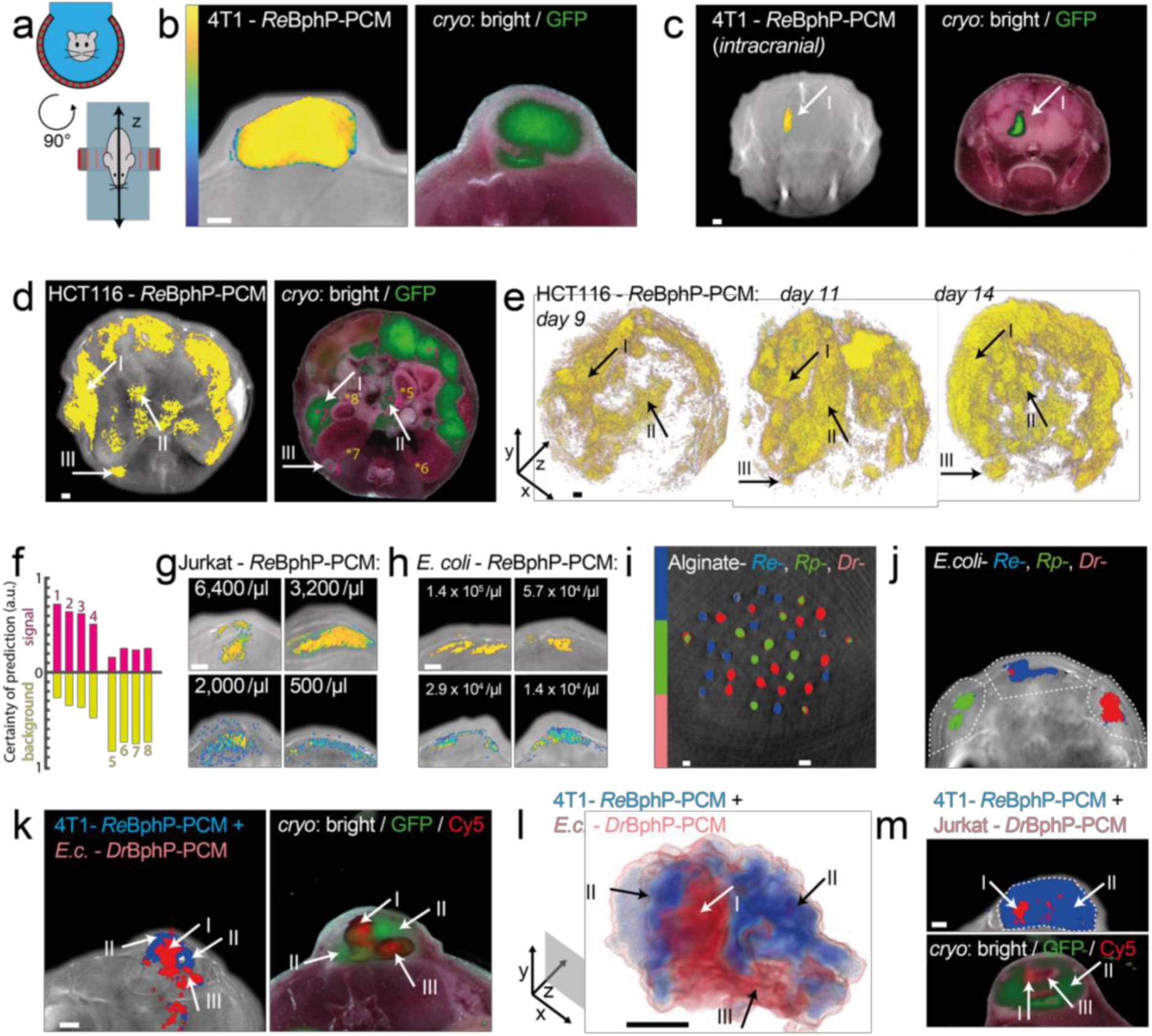
MSOT imaging of *Re*BphP-PCM and other rsOAPs. In certain experiments, GFP was coexpressed to allow fluorescence imaging of histology slices (excitation at 488 nm, emission at 510 nm). **a** Schematic of OA tomography used in this work. **b** 4T1 cells (0.8 x 10^6^ injected subcutaneously) stably expressing *Re*BphP-PCM and imaged on day 9. **c** 4T1 cells (0.7 x 10^6^ injected intracranially) stably expressing *Re*BphP-PCM imaged at a depth of 3.6 mm in the brain (Arrow I) immediately after injection. **d** HCT116 cells (1.5 x 10^6^ injected intraperitoneally) stably expressing *Re*BphP-PCM imaged on day 14. **e** Volume representation of the same mice at consecutive timepoints. (d and e) Arrows indicate distinctive tumor masses. **f** Certainty of prediction (weighted sum of tree scores) indicating quality of discerning label signal or background of ROIs shown in (d). **g** Imaging of the indicated concentrations of Jurkat T cells in Matrigel™ expressing *Re*BphP-PCM immediately after subcutaneous implantation; due to the polymerization process no homogeneity is expected. **h** Imaging of the indicated concentrations of *E. coli* expressing *Re*BphP-PCM in Matrigel™ immediately after subcutaneous implantation. **i** Imaging of an alginate bead phantom containing *E. coli* expressing rsOAPs *Re*BphP-PCM, *Rp*BphP-PCM, *Dr*BphP-PCM. **j** Imaging of *E. coli* (1.4 x 10^6^) expressing each of the three rsOAPs in Matrigel™ imaged immediately after subcutaneous implantation. **k** Imaging of a 4T1 tumor stably expressing *Re*BphP-PCM at day 9 (Arrow II and III) imaged immediately after *E. coli*(10^8^ cells) expressing *Dr*BphP-PCM have been injected into the tumor (Arrow I). Histology confirmation is inferred from fluorescence in *Dr*BphP-PCM (Cy5 only) and *Re*BphP-PCM (GFP primarily). **l** Volume representation of j. **m** Imaging of a similar 4T1 tumor expressing *Re*BphP-PCM (Arrow II) here injected with 10^6^ Jurkat T cells expressing *Dr*BphP-PCM (Arrow I; Arrow III refers to a necrotic patch). In panels (b-g), colormaps refer to R^2^ (detection quality); in (h-k), they indicate clusters showing distinguishable kinetics. All slices are single representative slices. All scalebars, 1 mm. Earlier timepoints and data from additional mice (panels b (n=3) and d (n=2)) can be found in Suppl. Figs. 7.

To assess the sensitivity of imaging with our rsOAPs, we imaged dorsal implants of Matrigel containing different numbers of Jurkat T lymphocytes stably co-expressing *Re*BphP-PCM and GFP in mice (Fig. 4g). We detected populations as small as 500 cells/μl, suggesting the potential for sensitive tracking of immune processes. Similarly, imaging of dorsal implants of Matrigel containing bacteria expressing *Re*BphP-PCM detected populations as small as 14,000 bacteria/μl (Fig. 4h). This sensitivity may be useful for studying and optimizing bacteria-based tumor therapies^21^.

A strong advantage of photo-controllable labels is the possibility to delineate multiple labels based on their individual switching kinetics. To demonstrate this, we imaged 1-mm alginate beads filled with purified *Re*BphP-PCM, *Rp* BphP-PCM or *Dr*BphP-PCM protein. All beads were unambiguously identified based on their switching kinetics (Fig. 4i). The same differentiation was achieved *in vivo* after implanting *E. coli* expressing each of the three rsOAPs into the back of mice (Fig. 4j).

Since the kinetics of photo-switching are energy-dependent, fluence changes due to light attenuation by surrounding absorbers – photochromic or static – complicates temporal multiplexing (Suppl. Note 5). Thus, one aim of our development of the fast switching *Re*BphP-PCM was to achieve a switching time constant clearly separate from other rsOAPs. We show that subcutaneously injected 4T1 cells expressing *Re*BphP-PCM are readily distinguished from infiltrating *Dr*BphP-PCM-expressing *E. coli* cells (intratumorally injected 108 bacteria Fig. 4k and l). This means that multiplexing is possible for co-registration studies, and that the concentrations of the labels can be estimated based on the convoluted kinetics (Suppl. Figure 8). Similarly, we show this for 4T1 and Jurkat T lymphocytes (intratumorally injected 106 cells Fig. 4m). Such, temporally unmixed multiplexed OA imaging of cells of the immune system enables following their function and involvement in disease mechanism *in vivo*, longitudinal on the organism level.

## Discussion

The combination of OA and transgenic rsOAP labels allows the tracking of specific cell populations *in vivo*, which can open up new possibilities for longitudinal studies of intact animals in diverse fields such as immunology, developmental biology, neurology, and cancer research. To support such studies, we describe next-generation rsOAPs that provide faster switching and greater resistance to photo-fatigue than existing rsOAPs, allowing highly sensitive detection, and importantly true multiplexing, without interference from hemoglobin or other abundant absorbers *in vivo*. These rsOAPs can be used with off-the-shelf equipment and our ML based open-access image processing code to detect populations of fewer than 500 cells *in vivo*. These tools will facilitate the wider use of OA imaging in life sciences, particularly for the study of cellular dynamics and interactions on the level of whole organisms.

## Materials and Methods

### Cloning

*Rp*BphP1^13^ was obtained from Addgene (Vladislav Verkhusha, plasmid # 79845). Mammalian optimized *Re*BphP was synthesized as gene strings (GeneArt, LifeTechnologies, Regensburg, Germany). All other BphPs used in the study have been a kind gift from Prof. Andreas Möglich, University of Bayreuth, Germany”.

For bacterial protein expression the coding sequences of all BphPs used in the study except *Rp*BphP1 were PCR amplified as a *NdeI/XhoI* fragment and cloned into the second multiple cloning site of pET-Duet1 vector (Novagene, Merck Millipore). *Rp*BphP1 was PCR amplified as a Nde/Pac1 fragment and cloned into the second multiple cloning site of pET-Duet1 vector Additionally, for biliverdin synthesis the heme oxygenase (HO) of *Nostoc sp*. was cloned by use of *NcoI/HindIII* into the first multiple cloning site of pET-Duet1.

For equimolar mammalian expression, first *Re*BphP1_P2A and mCherry were PCR amplified and then stitched using overlap PCR as an *EcoRI/XbaI* fragment and cloned in a pcDNA3.0 vector (Thermo Fisher). Later similar constructs for other BphPs were made by amplifying them as *EcoRI*/*NotI* fragment and inserted in place of *Re*BphP1-PCM in the above construct. The resulting plasmids allowed the equimolar co-expression of *Rp*BphP1, *Rp*BphP1-PCM, *Re*BphP-PCM, *Re*BphP-PCM or *Dr*BphP-PCM and mCherry proteins.

### Protein Expression and Purification

Proteins have been expressed in *Escherichia coli* strain BL21 (DE3) (NEB # C2527). In brief, plasmids expressing BphPs and HO were transformed into the BL21 host cells. Bacterial cells were grown in LB media supplemented with ampicillin at 37°C until the culture reached OD 0.6 followed by induction of protein expression by addition of IPTG and further incubation for 16 – 18 h at 22°C. Next day, the bacterial pellet was collected by centrifugation and pellet was resuspended in phosphate buffered saline (PBS). After cell lysis, proteins were purified by immobilized metal affinity chromatography (IMAC) in PBS, followed by gel-filtration on a HiLoad 26/600 Superdex 75pg (GE Life Sciences, Freiburg, Germany).

### Absorption and Fluorescence Spectroscopy

For absorption spectra the purification buffer was exchanged against phosphate-buffered saline and the proteins measured using a Shimadzu UV-1800 spectrophotometer (Shimadzu Inc., Kyoto, Japan) using a 100 μl quartz cuvette. To measure the ON (P_fr_) and OFF (Pr) spectra of respective proteins photo switching was carried out using 650/20 nm or 780/20 nm LEDs (Thorlabs) placed above the quartz cuvette in the spectrophotometer.

Fluorescence measurements for all BphPs were performed with a Cary Eclipse Fluorescence spectrophotometer (Varian Inc., Australia). Photo-switching was carried out as above. Fluorescence measurement was done by fixing excitation wavelength at 700 nm and emission wavelength at 720 nm. Excitation wavelength and emission slit were set to 5 nm and the absorbance at the excitation wavelength was always equal to 0.1 to avoid inner filter effects.

### Mammalian Cell culture

4T1 and Jurkat cells were maintained in RPMI-1640. HeLa cells and HCT116 cells were maintained in Dulbecco’s-modified Eagle’s medium (DMEM) and McCoy 5A medium respectively. All media were supplemented with 10% Fetal bovine serum (FBS) (Invitrogen) and antibiotics (100 U penicillin/ml and 100 mg streptomycin/ml). Cells were cultivated at 37° C and 5% CO_2_.

### Stable cell lines

#### Tissue culture

The Platinum-E and RD114 packaging cell lines were cultivated in cDMEM, HCT116 cell line was grown in McCoy (Life Technologies) and 4T1 and Jurkat cells were cultured in cRPMI. Alls media were supplemented with 10% FCS, 0.025% L-Glutamine, 0.1% HEPES, 0.001% gentamycin and 0.002% streptomycin.

#### Generation of constructs

*Re*BphP-PCM-IRES-GFP was amplified using specific primers (5’-ATTAGCGGCCGCGCCACCATGAGCGGCACCAGAG-3’, 5’-ATTAGAATTCTCACTTGTACAGCTCGTCCATGCCGTGAGTG-3’) and cloned into the mP71 using Not1 and EcoR1 restriction sites. The mP71 vector was a kind gift from Wolfgang Uckert.

#### Generation of cell lines

For retrovirus production, Platinum-E or RD114 packaging cells were transfected with the retroviral vector mP71-*Re*BphP-PCM-IRES-GFP using calcium phosphate precipitation. The supernatant of the packing cells was collected at 48 and 72h after transfection and purified from remaining cells by centrifugation at 1500 rpm at 4°C for 7 minutes. One day prior to transduction, non-tissue culture treated 48 well plates were coated with RetroNectin (Clontech) according to the manufacturer’s recommendations overnight at 4°C. After washing once with PBS, virus supernatant was added and centrifuged at 3000 g and 32°C for 2h. Virus supernatant was removed and cell lines (4T1, HCT116 and Jurkat) were added in 400μl of the respective medium supplemented with 1:100 Lentiboost Solution A, 1:100 Lentiboost Solution B (Sirion Biotech). Cells were then spinoculated at 800 g at 32°C for 1.5 h. After 5 days of culture, cells were sorted for high expression of GFP using flow cytometry.

### Mouse work

All animal experiments were approved by the government of Upper Bavaria and were carried out in accordance with the approved guidelines. For <u>4T1</u> xenografts of stably expressing *Re*Bphp-PCM and GFP, 0.8 x 10^6^ cells in PBS have been implanted in the back of FoxN1 nude mice (Charles River Laboratories, Boston, US) and maintained for 9 days. For <u>HCT116</u> cells expressing *Re*BphP-PCM and GFP, 1.5 x 10^6^ cells in 200 μl PBS have been injected intraperitoneally in FoxN1 nude mice and were maintained for 14 days. For <u>intra cranial injections</u> of stably expressing *Re*BphP-PCM_and GFP 4T1 cells, mice were first anesthetized according to the animal protocol. The head of the mouse was fixed in a Stereotaxic frame (David Kopf Instruments, Model 940), an incision in the skin was made using a scalpel and a small whole was drilled into the skull. Later, 5μl cells (0.14 x 10^6^ cells/μl) were injected slowly with a 10μl Hamilton syringe (26Gs). The incision in the skin was closed using Histoacryl^®^ (B. Braun Melsungen AG). The mice were scanned in MSOT and sacrificed immediately after scanning. For <u>Matrigel Implants</u> of Jurkat’s expressing *Re*BphP1-PCM, different concentrations of cells ranging from 8000 cells/μl to 800 cells/μl were implanted subcutaneously, in the back of the mice. Similarly, bacterial cells expressing *Re*BphP-PCM in different concentration (1.4 x 10^5^ to 1.4 x 104 cells/μl) were also implanted in the back of the mice. For multiplexing experiment, bacterial cells expressing rsOAPs individually with the concentration of 1.4 x 10^6^ cells/μl were implanted on the back of the mice in the same plane. For multiplexing experiment *in vivo,* <u>intra tumoral injections</u> bacterial cells expressing *Dr*BphP-PCM resuspended in PBS have been injected into the 4T1 tumor expressing *Re*BphP-PCM and GFP using an Insulin syringe with a 30G needle.

For all MSOT imaging mice have been anaesthetized using 2% Isofluran in O_2_. Anaesthetized mice were placed in the MSOT holder using ultrasound gel and water as coupling media. After termination of the experiments all mice have been sacrificed and stored at −80 °C for cryosectioning.

### MSOT setup and data acquisition

Phantom and mice data were acquired using a commercially available MSOT scanner (MSOT In Vision 256-TF, iThera Medical GmbH, Munich, Germany). In brief, nanosecond pulsed light was generated from a tunable optical parametric oscillator (OPO) laser and delivered to the sample through a ring-type fiber bundle. The wavelengths, 680 nm and 770 nm were used for photo-switching and imaging in phantoms as well as in mice. Light absorbed by the sample generates an acoustic signal that propagates through the sample and is detected outside the sample by a cylindrically focused 256-element transducer. The transducer array had a central frequency of 5 MHz (−6 dB was approximately 90%) with a radius of curvature of 40 mm and an angular coverage of 270°. Acoustic signals were detected as time-series pressure readouts at 2,030 discrete time points at 40 MS/s. The acquired acoustic data reconstructed using the ViewMSOT version 3.8.1.04 (iThera Medical GmbH, Munich, Germany) software with the following settings 50 kHz – 6.5 MHz, trim speed of 7.

## OA Data analysis

All data analysis was conducted using Matlab2018b. The data reconstructed with ViewMSOT was loaded into Matlab by iThera Matlab code (iThera Matlab, Version: msotlib_beta_rev75). All analyses were carried out with the code provided along with this manuscript (Suppl. Note 3). In brief: <u>Movement correction</u> was done by phase correlation preliminary to optimization-based image co-registration with the intensity and non-rigid co-registration of frames of the first cycle being used as reference. For further processing different features of the time series have been computed and are used for classification/switching label detection using a Machine learning model. For <u>fast Fourier transform</u> (fft) repetitive frequency of the whole concatenated signal for each image point are computed to identify signals corresponding to the illumination schedule. For <u>exponential fitting</u> the normalized mean kinetic of all cycles is used. Then the coefficients compared to an expected exponential kinetic are calculated and used as a quality measure. Here positive and negative exponential are considered. Using fit coefficients and quality of fit (R_2_) as measures only 77 % accuracy compared to a ground truth is achieved. Thus, additional features are invoked. Overall all analyzed features are: i ii) the coefficient for the exponential fit (exp(b(x+1)) and exp(b(x-1)+1) of the mean kinetic (mean of all cycles), iii) R_2_ of the fit, iv) the mean intensity over the concatenated signal, v) max-min of all the data at the pixel. vi – ix) median maximums and minimums of cycles along with standard deviation, x) number of cycles with positive or negative trend, xi) the length of the part of the cycle that shows a trend, i.e. at what point the signal vanishes in the noise, and xii) Fourier coefficient for the expected frequency defined by the photo-control schedule. All those are used as predictors values for an unmixing model based on <u>random forest approaches</u>^20,22^ – for overall model, trained on 4T1 day 9 as well as highest concentration Jurkats T lymphocytes. We used 50 trees in the ensemble as further increase of number did not lead to out of bag error decrease. This approach resulted in model performance increased to 96% of positive predictive value for ground truth. (See Suppl. Note 4 for more details on the use of ML in this work).

For visualization data was not further processed and is shown against the respective slice at 680 nm as anatomy information, except in the case of 4T1 injected in brain where the anatomy is shown at 900 nm. Representative slices are shown.

For clustering appropriate ranges of the kinetic parameter where chosen on the unmixed data to distinguish different labels.

## Cryosectioning

After sacrificing the mice were cryopreserved at −80 °C. To detect the fluorescence in tumors, the respective part of the mouse was embedded in Tissue-Tek^®^ O.C.T. (TM) (Sakura Finitek Europe B. V., Zoeterwonde, Netherlands). 10 μm sections were cut (Leica CM1950, Leica Microsystems, Wetzlar Germany) for brain, 4T1, HCT116 mice at the interval of 150 μm, 250 μm and 500 μm respectively and imaged using a 482/35 nm bandpass for excitation and 535/38 nm bandpass filter for detection of GFP fluorescence. Images were taken using an Andor LucaR CCD camera (DL-604M, Andor Technology, Belfast, UK) with 10 s exposure and a gain of 10.

## Bacterial Cell immobilization by entrapment and MSOT Imaging

A 2-4% (*w*/*v*) aqueous solution of sodium alginate was prepared in PBS. *Escherichia coli* strain BL21 cells expressing rsOAPs were harvested by centrifugation (4000 rpm, 20 min) and resuspended in PBS. The cell suspensions were then mixed with sterile alginate. Beads were formed by filling the alginate-cell mixtures in the syringe with 30G needle followed by centrifugation at 300 rpm which allowed the addition of the mixtures into sterile CaCl2 (200 mM). The cell-containing beads, 1 mm in diameter, were allowed to solidify for 10 min before CaCl2 was replaced by fresh distilled water. The cell beads were then randomly distributed in the agar phantom with 1.5% (w/w) agar and 3.5% (v/v) intralipid emulsion and imaged in MSOT as described elsewhere.

## Optoacoustic characterization of proteins

For optoacoustic characterization of rsOAPs, custom-made experimental set-up was used as described earlier 19. Briefly, Nanosecond excitation pulses were generated by an optical parametric oscillator (OPO) laser (Spitlight-DPSS 250 ZHGOPO, InnoLas) running at a repetition rate of 50 Hz Constant pulse energy was ensured by use of a half-wave plate in a motorized rotation stage (PRM1Z8, Thorlabs) and a polarizing beam splitter; using a lookup table and adapting the polarization with the half-wave plate, we kept the power constant at ~1.3 mJ (otherwise mentioned) over the whole illumination schedule. Samples were injected into an acoustically coupled flow chip (μ-Slide I 0.2 Luer, hydrophobic, uncoated, IBIDI) and illuminated from one side by use of a fiber bundle (CeramOptec) at a constant pulse energy of ~1.3 mJ at the fiber output. Photo-switching was carried out by illuminating the sample alternatively with 650 nm and 780 nm light. Optoacoustic signals were detected with a cylindrically focused single element transducer (V382-SU, 3.5 MHz, Olympus) followed by signal were amplification by 60 dB via a wide-band voltage amplifier (DHPVA-100, Femto) and digitized at 100 MS/s with a data acquisition card (RZE-002 400, GaGe). Dependency of P_fr_→Pr conversion on 770 nm pulse energy was measured with different pulse energies (0.4 mJ, 0.7 mJ, 1.0 mJ and 1.3 mJ). Dependency of P_fr_→Pr conversion on repetition rate of laser was measured with three different laser repetition rates (10, 25, and 50 Hz). Effect of different switching ON wavelength and resulting dynamic range at 770nm was measured using different switching ON wavelength ranging from 630 nm to 680 nm.

## Supplementary Materials

Suppl. Table 1: Overview of native BphPs and truncated versions

Suppl. Fig. 1: Absorption spectra and OA photo-switching of *Rp*BphP1

Suppl. Note 1: Truncation strategy

Suppl. Fig. 2: Monomeric nature of the two developed rsOAPs

Suppl. Fig. 3: Mammalian expression of rsOAPs

Supp. Fig. 4: Power and Repetition rate dependency of rsOAPs.

Suppl. Fig. 5: Switching wavelength dependence of *△OA_On___O_ff* for *Re*BphP-PCM. Suppl. Fig. 6: Information content analysis.

Suppl. Note 2: Information content analysis

Suppl. Note 3: Details on the analysis and script availability and functionality

Suppl. Note 4: Machine learning (ML)

Suppl. Fig. 7: Representative images of all mice

Suppl. Note 5: Challenges of unmixing entangled populations

Suppl. Fig. 8: Concentration dependent change of switching kinetics.

Suppl. Fig. 9: Dependence of switching kinetics on depth in switchable material

## Acknowledgments

The authors wish to thank Prof. Andreas Möglich for providing the wildtype BphPs, Ruth Hillermann for technical assistance and Armando C. Rodríguez for discussions on the manuscript.

## Funding

K.M. and A.C.S receives funding from DFG (STI656/1-1). V.R.B. receives funding from DFG SFB 1054 (TP B09)

## Author contributions

K.M. performed all measurements. For *in vitro* data with help from J.P. F.-W.; For *in vivo* measurements with assistance from U.K. and occasional help from V.G.; K.M. analyzed all *in vitro* data. All *in vivo* data was analyzed by M.S. along with K.M.; Analysis code and GUI written by M.S.; S.G. established stable cell lines; A.C.S. wrote the manuscript and conceived the project together with K.M. and M.S.. V.R.B. and V.N. contributed to the manuscript.

## Competing interests

V.N. is a shareholder of iThera Medical GmbH. All other authors declare no competing interests.

## Data and materials availability

### Data Availability

A reduced source data contents file is available online. Due to size limitations of depositing raw imaging data the data that support the findings of this study are available from the corresponding author upon request.

### Code Availability

Detailed code is available from the corresponding author upon request. A version of the code meant for public use together with a GUI can be found at https://gitlab.lrz.de/ga45huk/rsoap_analysis/.

## Notes

### Competing Interest Statement

VN is a shareholder of iThera Medical

