## Supplementary material for "Multiplexed whole animal imaging with reversibly switchable optoacoustic proteins": suppl

**Suppl. Table 1: Overview of native BphPs and truncated versions (PCM)** screened in this study for favorable characteristics as rsOAPs.

| Protein | Photoswitching state | Absorption maxima (nm) | Molar absorption coefficient ($M^{-1}$ $cm^{-1}$) | Emission maxima (nm) | QY[a] | switching mode | OA switching dynamic range (Pfr/Pr) at 770nm [b] | photoacoustic generation efficiency (PGE) [c] | switch-off halftime $t_{1/2}^{off}$ (s) [d] | switch-on halftime $t_{1/2}^{off}$ (s) [d] |
|---|---|---|---|---|---|---|---|---|---|---|
| *Rhizobium etli* (*Re* BphP1-PCM) | Pfr(ON) | 708 / 760 | 92287 / 84520 | none | 2.94 | neg | 4.821 ± 0.254 | 1,12 ± 0,02697 | 0,1815 | 0,2441 |
| | Pr(OFF) | 704 | 11272 | 720 | | | | | | |
| *Deinococcus radiodurans* (*Dr* BphP1-PCM) | Pfr(ON) | 750 | 87030 | none | 2.90 | neg | 2.871 ± 0.018 | 1,074 ± 0,01722 | 0,7045 | 0,2406 |
| | Pr(OFF) | 700 | 98246 | 720 | | | | | | |
| *Rhodopseudomonas palustris* (*Rp* BphP1-PCM) | Pfr(ON) | 758 | 74812 | none | 1.84 | neg | 2.228 ± 0.143 | 0,8301 ± 0,06495 | 0,4202 | 0,2851 |
| | Pr(OFF) | 688 | 41197 | 720 | | | | | | |
| *Rhodopseudomonas palustris* (*Rp* BphP1) | Pfr(ON) | 756 | 78300 | none | none | neg | n.d | n.d | 0.65 [e] | n.d |
| | Pr(OFF) | 678 | 87500 | | | | | | | |
| *Azorhizobium caulinodan* (*Ac* BphP-PCM) | Pfr(ON) | 702 / 750 | 56744 / 46427 | n.d | n.d | neg | n.d | n.d | n.d | n.d |
| | Pr(OFF) | 699 | 50836 | | | | | | | |
| *Xanthomonas campestris* (*Xc* BphP-PCM) | Pfr(ON) | 682 / 750 | 51993 / 30091 | n.d | n.d | neg | n.d | n.d | n.d | n.d |
| | Pr(OFF) | 682 | 53666 | | | | | | | |
| *Agrobacterium tumefaciens* (*At* BphP2-PCM) | Pfr(ON) | 756 | 50663 | n.d | n.d | neg | n.d | n.d | n.d | n.d |
| | Pr(OFF) | 756 | 46829 | | | | | | | |
| *Agrobacterium vitis* (*Av* BphP-PCM) | Pfr(ON) | 755 / 709 | 51219 / 48137 | n.d | n.d | neg | n.d | n.d | n.d | n.d |
| | Pr(OFF) | 702 | 48468 | | | | | | | |
| *Pseudomonas syringae* (*Ps* BPhP2-PCM) | Pfr(ON) | 708 / 750 | 57042 / 50540 | n.d | n.d | neg | n.d | n.d | n.d | n.d |
| | Pr(OFF) | 702 | 72999 | | | | | | | |

[a] QY was determined with Abs equal to 0.1 at $\lambda$ex = 680nm, [b] Data shown is from MSOT where 680/770nm was used for photo-switching and dynamic range is defined as the ratio of OA signal Intensity between the Pfr and Pfr upon illumination with 770nm, [c,d] Pulsed illumination (1.3 mJ/pulse, 7 ns pulse length, 50 Hz repetition rate), & 650/770nm was used for photo-switching, [e] Ref19. ±, represent standard deviation. n.d, data not defined in this study.

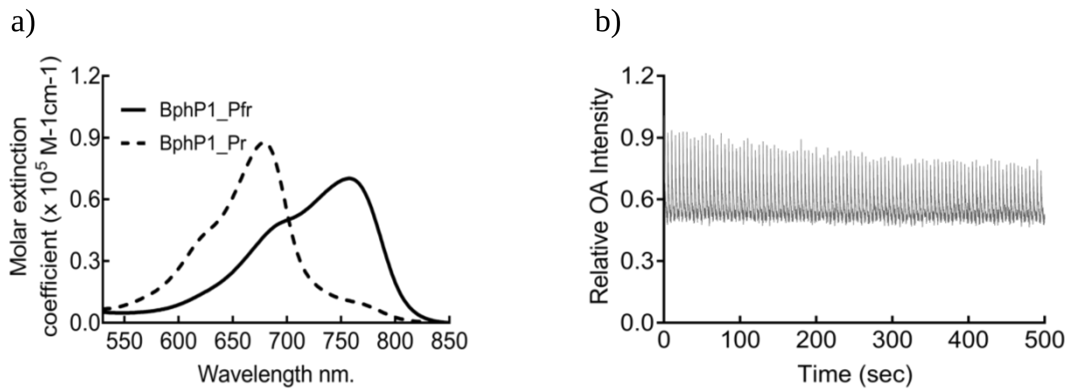


**Suppl. Fig. 1: Absorption spectra and OA photo-switching of *Rp*BphP1, parent of *Rp*BphP1-PCM. a** Absorption spectra of Pfr (solid) or Pfr (dotted) state, measured after illumination of the purified protein with a 650/20 nm or 780/20 nm LED placed above the cuvette, respectively. **b** OA-switching by 50 pulses of 770 nm light per cycle (Pfr → Pr). The back-transition induced by 50 pulses of 680 nm light per cycle (Pr → Pfr) is omitted for clarity.

**Suppl. Note 1: Truncation strategy and molecular explanation for distinct switching speed of rsOAPs in this study.**

Towards identifying a BphP that shows favourable switching characteristics and monomeric behaviour when truncated to the minimum domains necessary for photo-switching (PAS, GAF, PHY domains, PCM = photosensory core module) eight BphPs were truncated or obtained as truncated version and tested for photophysical characteristics favourable for OA imaging (high absorption coefficient and high change in signal upon photo-switching at 770 nm, Suppl. Table 1). Among the screened BphPs, the truncation of *Rhizobium etli* BphP (*Re*BphP-503) was the one showing the highest absorptivity and strongest change in signal at 770 nm upon photo-switching.

In a next step we explored if slightly different truncation sides might further improve the photophysical characteristics and since no structural information was available for *Re*BphP we modelled the protein structure using I-TASSER (reference model: 6g1y). Based on this data we choose additional truncation points at position 524 and 547 which according to the modelled structure contain the minimum PAS-GAF-PHY photosensory core domains (503) together with extra amino acids from the annotated linker between PHY and Histidine Kinase domains. We choose the 524-length version (hereafter named *Re*BphP-PCM) because it shows equal switching behaviour to the longer 547 and wildtype but is superior to the shortest truncation of 503 amino acids (Suppl. Figure 2a and b). The loss in signal change upon photo-switching of the shortest version (*Re*BphP-503) might be attributed to the PHY domain capping loop not completely shielding the chromophore during photo-switching.

For *Rp*BphP1 we followed a similar truncation strategy with truncation at positions 589, 554 and 522. Interestingly, in this case the 522 amino acid version (hereafter named *Re*BphP-PCM) showed a switching behaviour most similar to the full-length protein while the longer versions showed lower signal changes after photo-switching (Suppl. Figure 2c and d). In earlier works it was suggested that a truncation of *Rp*BphP1 is not producing switchable variants due to the loss of HOS domain [1]. We could not find such a relationship and yielded a monomeric protein after truncation at position 522 (Suppl. Fig. 4) showing full wildtype switching behaviour (Suppl. Fig. 2 and 3).

The three BphP-PCMs used in this study show distinctive switching speeds which is the reason for our ability to discriminate the proteins *in vivo* successfully to facilitate true multiplexing. Based on existing crystal structures of *Rp*BphP1 (4gw9) and *Dr*BphP (4o01) and our model for *Re*BphP-PCM we analyzed the molecular determinants of the different switching speed, primarily the metastability of the $P_{fr}$ state. As already pointed out this is primarily influenced by the PHY loop capping the BV binding pocket [2–4]. In *Rp*BphP1 and *Dr*BphP the D-ring of the BV chromophore is primarily stabilized via an interaction with D201 and D207, respectively. The primary interaction with the capping loop of the PHY domain is via S468 in both proteins (Suppl. Figure 1a). In contrast, in the structure of e.g. *Pa*BphP (3c2w) we find D194 interacting with R453 additionally to the aforementioned serine, here S459. Probably this second ionic interaction in the opposite direction weakens the stabilization of the $P_{fr}$ conformer of the chromophore (Suppl. Figure 1b). Sequence comparison and our homology model suggests the same scenario for *Re*BphP-PCM suggesting that a less stabilized $P_{fr}$ state might favor the 770 nm induced transition from $P_{fr}$ to $P_r$ and reduce the efficiency of the dark relaxation form $P_r$ to $P_{fr}$ thus result in faster switching.

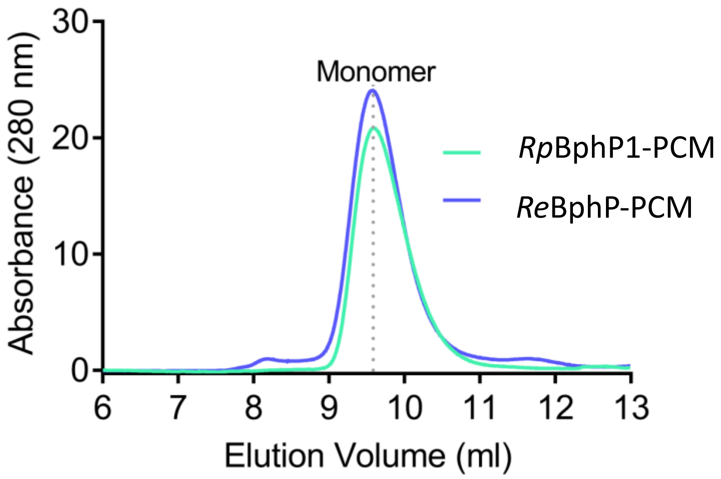


**Suppl. Fig. 2: Monomeric nature (~60 kDa) of the two developed rsOAPs.** The purified proteins in phosphate buffered saline were run on an analytical size exclusion chromatography (Superdex 75, 10/300 GL (GE Life Sciences, Freiburg, Germany)) at 0.8ml/min.

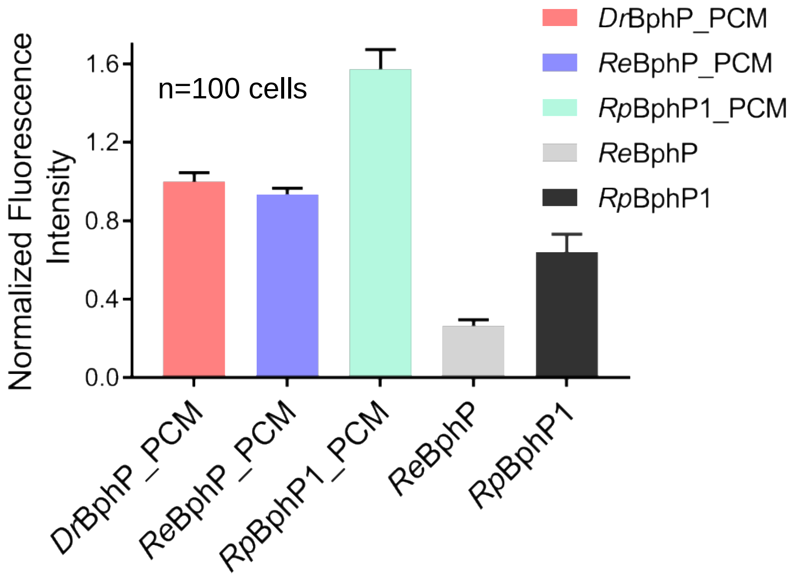


**Suppl. Fig. 3: Mammalian expression of rsOAPs.** The respective rsOAPs-P2A-mCherry constructs were transfected in HeLa cells using Lipofectamine-2000 using manufacture's protocol. Cells were visualized after 48 hours using a fluorescence microscope with Exc: 570/20 nm and Em: 710/20 nm. For quantification, 100 cells for each rsOAPs were selected and the fluorescence integrated density was calculated using ImageJ. Bar graphs represent the mean fluorescence intensity normalized by cell area. For convenience the data is further normalized to the value of *Dr*BphP-PCM for easy comparison. Error bars represent the standard deviation (n= 100 cells).

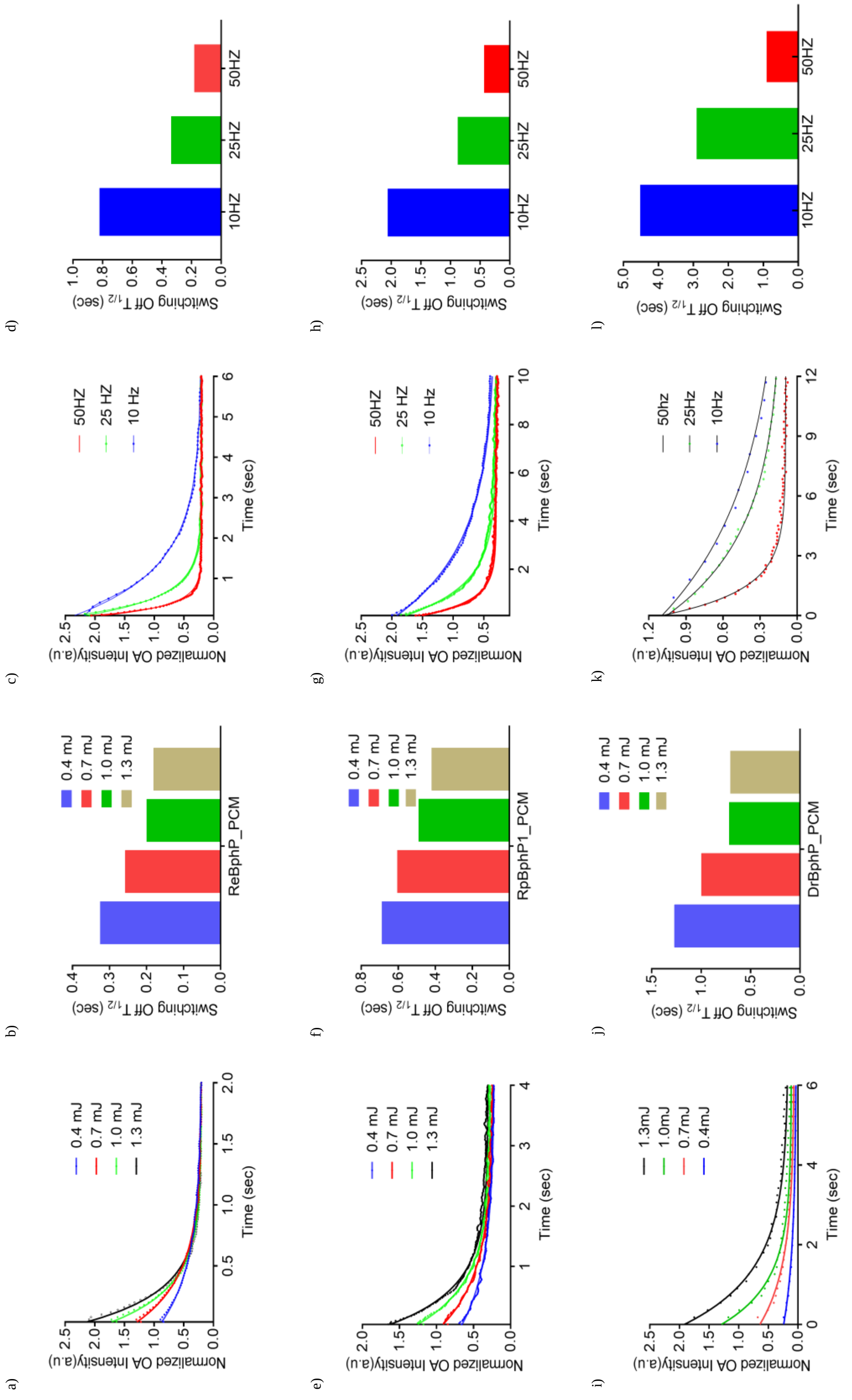


**Suppl. Fig. 4: Power and Repetition rate dependency of rsOAPs. a-d** *Re*BphP-PCM, **e-h** *Rp*BphP1-PCM and **i-l** *Dr*BphP-PCM. All data was recorded using an analytical OA spectrophotometer. Details can be found in the method section.

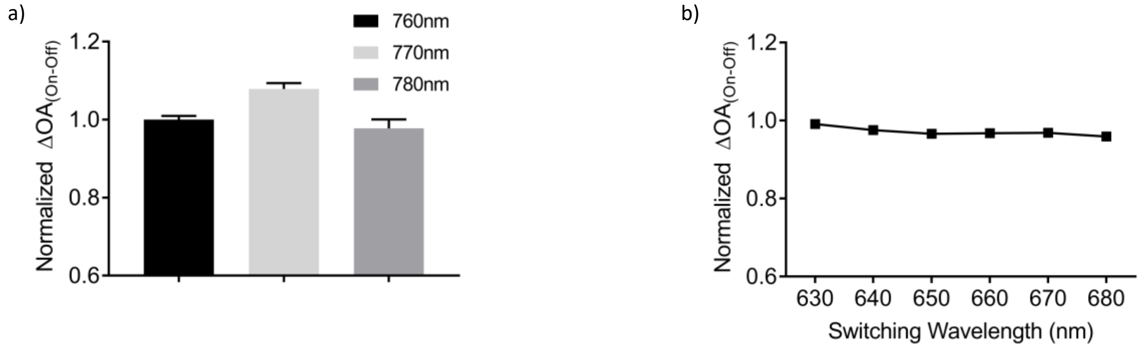


**Suppl. Fig. 5: Switching wavelength dependence of $\Delta OA_{On-Off}$ for *Re*BphP-PCM. a** Normalized $\Delta OA_{On-Off}$ after off-switching with 760, 770 or 780 nm after using 680 nm for photo-switching to the on-state. **b** Normalized $\Delta OA_{On-Off}$ after off-switching with 770 nm in relation to different on-switching wavelengths used (630 – 680 nm). Dependence on ON-switching wavelength was measured with an OA analytical spectrometer instead of MSOT, see methods for details.

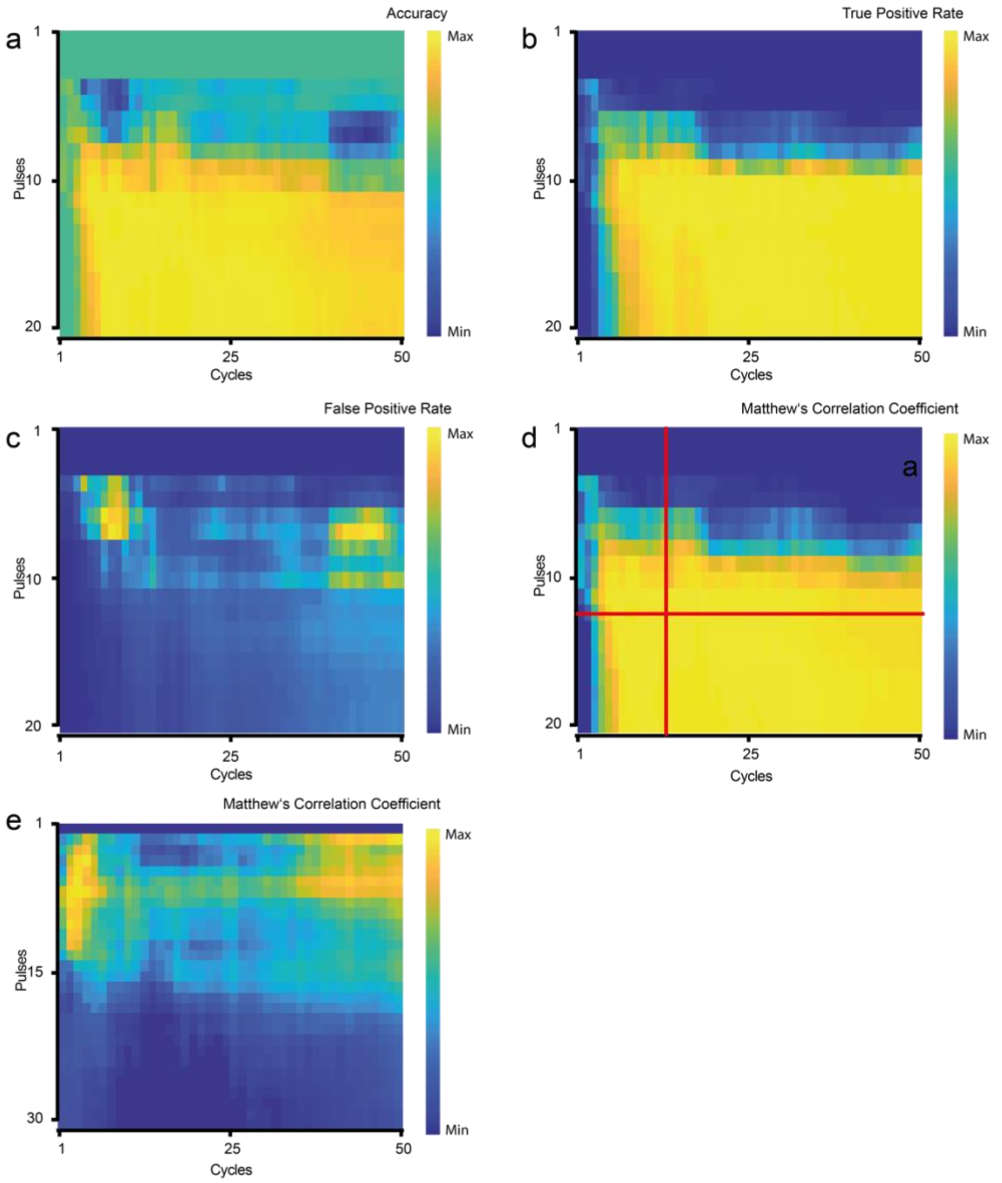


**Suppl. Fig. 6: Information content analysis.** Shown are the accuracy (**a**), true positive rate (**b**) and false positive rate (**c**) as well as the Matthew's correlation coefficient (**d**) of a dataset (4T1 mouse bacterial injection pre-scan) analyzed for combinations of number of cycles and pulses per cycle, respectively. The ground truth was established by comparing the expected region against histology. The red lines in (d) represent the number of cycles or pulses expected to give a good result with no significant improvement if more pulses or cycles are used. For this calculation the maximum number of pulses and cycles is used as best scenario and then reduced until changes in the number of detected pixels occur. This measure correlates very good with the observations from the info content analysis and allows to give an estimate on the minimum of cycles and pulses needed for a given imaging situation (e.g. a longitudinal imaging approach) without requiring histology. We note that the patches of high numbers of false positives at ~5 pulses / ~10 cycles and ~8 pulses / ~45 cycles, respectively, result from bad quality of the respective cycles in the analyzed dataset. **e** Matthew's correlation coefficient for a dataset with comparably low signal vs. noise. Here more pulses are show to visualize the impact of longer recording windows on the information content if noise is high. Further, the stretch between 10 and 30 cycles visualizes the impact of "bad" cycles, e.g. due to mouse movement or other artifacts.

**Suppl. Note 2: Information content analysis (Suppl. Figure 6).**

The temporal unmixing is based on analyzing the repetitive time varying patterns of the photo-switchable label in dependence on the applied photo-control schedule (i.e. the number of light pulses per wavelength per modulation cycle). Consequently, the number of pulses dictates how much the label population is switched to the other state, i.e. how much time varying kinetic can be analyzed per cycle or how much surplus data (noise) is recorded per cycle when the protein is already completely off. Especially the latter is observed to strongly reduce automatic detection quality (Suppl. Fig. 8e). Beyond that sometimes higher numbers of pulses might be necessary if the switching kinetics time window correlates with other periodic signal like breathing or heartbeat which can add a periodically varying signal to the data and can be extracted by having a longer recording window per photo-switching cycle (i.e. more pulses).
The number of cycles determines how many repetitions are available for the subsequent analysis to differentiate the label from background. I.e. more cycles result in better denoising. By analyzing combinations of number of cycles and pulses per cycle and the resulting precision in verifying the ground truth (histology) we could show that already small numbers of cycles and pulses are enough to produce reliable results (Suppl. Figure 8). Further, by a converging algorism we could determine those minimal numbers of pulses and cycles without the ground truth of histology (see Suppl. Figure 8 for details). Such analysis is clearly dependent on the sample in regard to label density, depth of label in the mouse body and heterogeneity of labeled structures. However, it can be valuable when conducting time-resolved analysis by determining the minimal possible image dwell-time with the first measurement and then setting the parameters in further measurements accordingly. For example, for a small 4T1 tumor at the begin of its growth we see no improvement of Matthews' correlation coefficient after 11 pulses with 7 cycles. For our 10 Hz system that would result in a dwell time per tomography slice of ~8s. On the protein level, much faster switching is achievable thus it can be envisioned that with faster systems (higher laser repetition rate) a time resolution < 1 s is easily achievable.

**Suppl. Note 3: Details on the analysis and script availability and functionality**

Beyond the temporal-development of the signal at each pixel the data carries a multitude of additional information. The selection how this signifies the presence of a switchable label is not straight-forward. Accordingly, we choose to employ models derived from classic machine learning (ML) to analyze the data (see Suppl. Note 4).

In the following the data analysis is explained in detail (Fig. 3). All aspects of data analysis and training of ML models are part of the code freely accessible under the GNU public license at https://gitlab.lrz.de/ga45huk/rsoap_analysis/. Assistance using the functions and the unmixing strategies can be obtained from the authors.

***Data calculation function (calculation.m)***

As an input the code requires a Matlab array with the consecutive time-varying data per pixel. E.g. a 300 x 300 pixel image recorded with 25 cycles and 10 pulses of e.g. 680 nm ($WL_A$) and 10 pulses of 770 nm light ($WL_B$) would yield a 300 x 300 x 500 array with the last dimension being: $(p_{680\_1} \dots p_{680\_10}, p_{770\_1} \dots p_{770\_10})_{cycle\_1} \dots (p_{680\_1} \dots p_{680\_10}, p_{770\_1} \dots p_{770\_10})_{cycle\_25}$

Such arrays can be easily extracted from the instrument software. In this work, using the In Vision 256-TF MSOT (iThera Medical, Germany), the code for extracting the data in Matlab format can be obtained from the manufacturer's website (ithera-medical.com, ithera_data.mat contains an example script for this task). But any other OA instrument with a tunable laser source in the NIR range would be equally suitable, e.g. the Vevo LAZR-X (FUJIFILM VisualSonics, Canada), the TriTom system (Photosound, U.S.A.) or the LOIS-3D (TomoWave, U.S.A.).

Beyond this input array the main script requires the photo-control schedule details (e.g. 25 cycles of 10 pulses $WL_A$ and 10 pulses $WL_B$), which wavelengths to process and a number of optional parameters, regarding selection of a region of interest, the use of fluence and motion correction and the use and characteristics of initial coarse-fitting. Beyond the wavelength and illumination schedule all parameters are just relevant for reducing computation time and can be left in the default state. The function returns a Matlab file with all features for each pixel of the given dataset. The features are: i-ii) the coefficient for the exponential fit (exp(b(x+1)) and -exp(b(x-1)+1) of the mean kinetic (mean of all cycles), iii) $R^2$ of the fit, iv) the mean intensity over the signal, v) amplitude (max-min) of all the signal, vi-ix) median maximums and minimums of cycles along with standard deviation, x) number of cycles with increasing or decreasing trend, xi) the length of the part of the cycle that shows a trend, i.e. at what point the signal vanishes in the noise, and xii) Fourier coefficient for the expected frequency defined by the photo-control schedule.

***Data analysis function GUI***

This function and the associated GUI analyze the data with various parameters that can be user selected. It is much faster than the initial step of "data calculation" and allows the testing of different parameter sets. In default mode this function just requires the calculated data from the previous function as well as a pre-calculated classification model. The overall classification model used for this work, covering superficial and deep-seated tumors as well as small cell populations with our labels can be found along with the code (features.mat). All parameter ranges can be also set manually; note that this results in preselection by the changed parameter ranges and omitting those features in the model. In the GUI the features are order by their predictor importance evaluated by model performance on validation dataset within training for the overall model used in this work. New classification models based on different training data can be generated by the user, see below.

The script returns Matlab matrices of the unmixed image in the following formats: coefficients of the exponential fit (useful to distinguish multiple labels) and $R^2$ of the exponential fit (a criterion for the quality of detection). Both data are also returned as an .png image together with overlays with an anatomy image at the specified wavelength (set by anatomy_WL parameter). Finally, a Matlab structure as well as text file with statistics overview, main values and unmixing summary are outputted.
Setting the information content flag additionally provides information after how many cycles and pulses no significant changes in the result could be detected. Allowing to adjust parameters of the illumination schedule (cycles, pulses) for subsequent measurements of comparable samples and thus lower dwell time (Suppl. Note 2).

**Suppl. Note 4: Machine learning (ML) approaches and model training (training.m)**
In principle the detection of rsOAPs relies on detecting a periodic exponential behavior defined by the given photo-control schedule. However, we found that beyond analyzing just this periodicity and the slope of the exponential change it proofed to be useful to incorporate a number of other features derived from the data (see Suppl. Note 3). The direct involvement of the two former is straight forward and dictated by the photo-control schedule; while however, defining ranges for the other features was not directly deducible. Thus, we resorted to the use of classic machine learning (ML) approaches with the complete set of features being: i-ii) the coefficient for the exponential fit (exp(b(x+1)) and -exp(b(x-1)+1) of the mean kinetic (mean of all cycles), iii) $R^2$ of the fit, iv) the mean intensity over the signal, v) amplitude (max-min) of all the signal, vi-ix) median maximums and minimums of cycles along with standard deviation, x) number of cycles with increasing or decreasing trend, xi) the length of the part of the cycle that shows a trend, i.e. at what point the signal vanishes in the noise, and xii) Fourier coefficient for the expected frequency defined by the photo-control schedule.

In order to incorporate those additional features in the detection algorithm we implemented two supervised machine learning techniques typically used for classification problems. Those are decision tree-based learning (random forest) [5] and support vector machine (SVM) methods [6]. In classification two labels were used: background and signal. For the training and validation dataset labeling was done according to ground truth acquired from histology. Training and validation datasets were built from datasets of 4T1 day 9 (Suppl. Figure 7c and d) as well as highest concentration Jurkats T lymphocytes (Figure 4g). As in those datasets we observe generally large portions of background compared to smaller regions with label, in order to eliminate redundant background information, the amount of signal and background labeled data points were balanced by excluding background points using mirror-reachable rule [7]. The remaining points were randomly divided between training and validation datasets in relation of 60% to 40%. As predictors the above-mentioned set of features was used in both models.

*Random forest*
For reducing variance of the decision trees a bagging technique was used [8]. The number of trees in the forest was set to 50, as a further increase of the number of trees did not lead to a decrease of the out-of-bag classification error, which is less than 0.04 for the final model. Reduction of the number of features was tried but led to accuracy loss. Similarly, predictor association estimation, which shows correlation for each pair of features, was less than 0.3, so that none of the features can be aggregated or disregarded. The resulting model shows 96.6% accuracy on the validation dataset.

*Support Vector machine (SVM)*
As we have a multidimensional feature space with complicated feature relationships, a non-linear SVM with a Gaussian radial basis function kernel was used [9]. Such a model showed even higher accuracy on alike datasets than the random forest described above, when trained for a single dataset (only 4t1 or only Jurkat); accuracy on validation dataset here is 98.9%. However, when trained on a combination of different datasets (4T1, Jurkat) the models start to overfit.

We tested both approaches on independent datasets which were not used in training and are of a different type than the training data (HCT116 scans together with the respective histology as ground truth). Here, the decision tree-based model showed a clear advantage with a smaller number of false positives. This suggest that random forest models might be better suited for unknown data of a different type while SVM could be better suited in repetitive imaging experiments delineating labels in relatively same depth and density.

***Model training (training.m)***

Along with the code we provide the overall general model described above based on the bagged tree approach that was used throughout this work. However, given a more expanded imaging project with ground truth available from histology of initial scans we encourage the user to build their own models using the training script. In the GUI the user is asked for each training data set to determine the analysis ROI (mostly the mouse body region) as well as the label location as informed by ground truth (e.g. histology). Further the model type bagged tree or support vector machine has to be set. The script returns the confusion matrix, ROC curve and value range of main features in graphical representation for analysis of the training. Along with that the model is saved as a .mat for use in the analysis script (Suppl. Note 3).

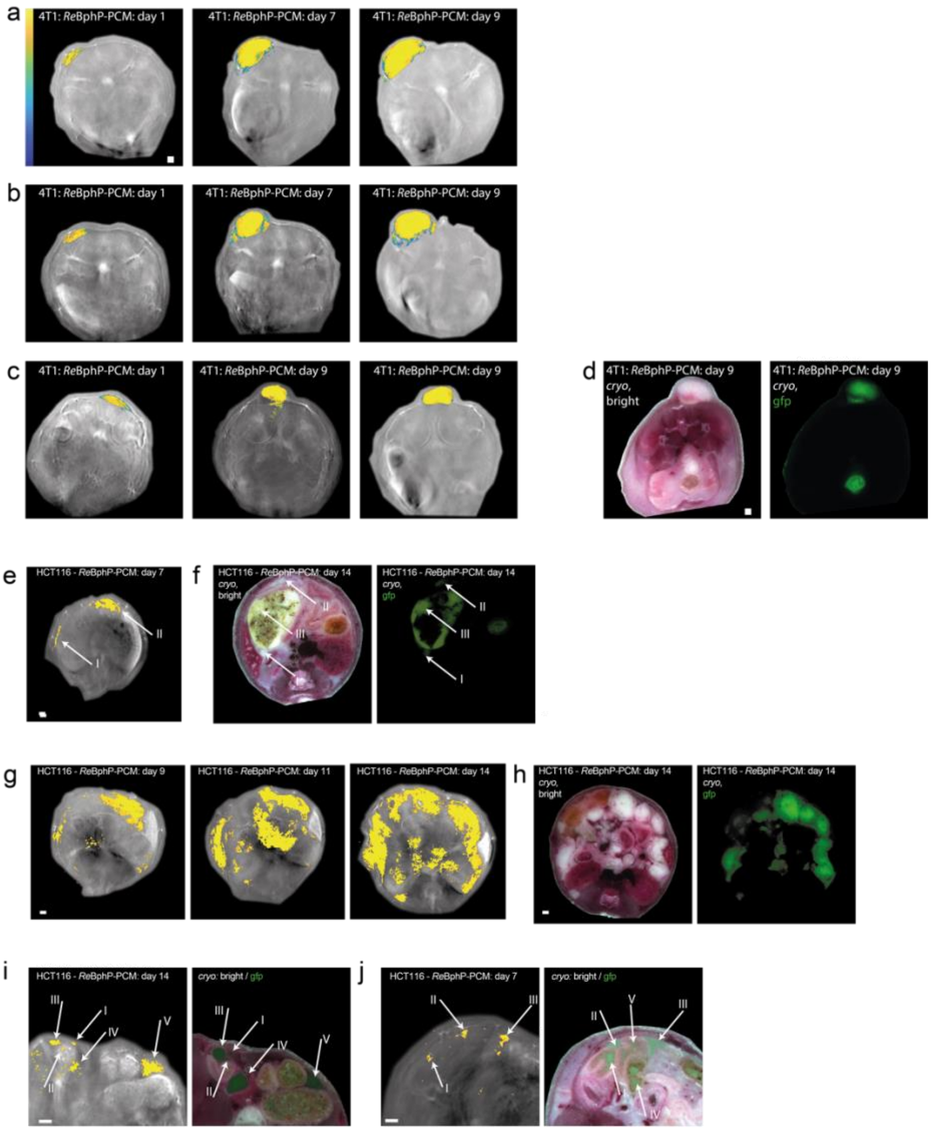


**Suppl. Fig. 7: a-c** Representative images of all mice bearing 4T1 tumors expressing *Re*BphP-PCM (n=3) at selected timepoints. Image shown here are colored in $R^2$ indicating the fit quality of the mean kinetic to the exponential model, i.e. the quality (or certainty) of the unmixing for the given pixel. **d** Cryo-slices imaged brightfield (left) and GFP (right) for the mouse in (c) at day 9. Mice bearing HCT116 tumors expressing *Re*BphP-PCM at selected timepoints. **e,g** Representative OA images of all HCT116 mice (n=2). **f,h** Histology as brightfield and GFP-fluorescence at the last timepoint after sacrificing the animal. For (e and g) tumor regions are marked (Arrows I and II) to distinguish from intestine autofluorescence (Arrow III). **i,j** Detail of a different slice of the HCT116 mouse, day 14 from (f and h) as well as day 7 from (a and b) shown together with the corresponding histology. Several tumor masses can be clearly associated with histology (arrows I to V in i and I to III in j). The smallest tumors (arrow I and II in i and I in j) roughly have a size of 250 µm$^2$ corresponding to < 10,000 HCT116 cells. We note that due to contributions from OA signals outside of the primary z-plane we see additional signals e.g. on the left part of (i). Furthermore, intestine can contribute to false-positive GFP-fluorescence (arrow IV in j). Finally, we point out that very low numbers of disperse tumor cells as apparent by the light GFP-fluorescence (arrow V in j) are not adequately picked up here.

**Suppl. Note 5: Challenges of unmixing entangled populations based on temporal unmixing**

The photo-switching process is positively correlated with the light fluence. Thus, the switching kinetics depend on how much lights effectively reaches the proteins in the given voxel [10]. In tomographic whole animal imaging the light fluence is dependent on the depth, wavelength and the absorbance nature of the tissue in the light path. This leads to an attenuation of the light fluence in deeper layers which can be roughly estimated by light fluence models [11]. For larger structures expressing photochromic labels, i.e. the light travels through photo-switching absorbers before it reaches the voxel under consideration, this is even further complicated by the photochromism leading to a change in absorbance with a kinetic dependent on the label and the light fluence. Thus, the switching kinetic of such a voxel is a convolution of its individual switching kinetic and the kinetic of the light fluence change of the above layers of switchable material (Suppl. Figure S9a-d). However, we observe this effect to be relatively neglectable in smaller structures and small compared to heterogeneities anyways present in the area (Suppl. Figure S9e-h)

Additionally, due to this effect non-switchable absorbers beneath switchable labels can appear switching due to their effective light fluence changing with the characteristic time kinetics of the above absorbers. For negative switchers [12] this effect leads to a positive kinetic which can be easily separated from the negative kinetics of the label itself. For unmixing of multiple labels of different types, where label A lies in the light path to label B, this results in the resultant kinetic being a convolution of the photo-switching kinetics of label A and B.

The temporal unmixing of multiple labels based on their, intrinsic, different photo-switching characteristics relies on performing an exponential fit on the kinetic of the mean over multiple switching cycles for each pixel and based on the resulting fit coefficients discriminate among the different labels. Due to the entanglement of the kinetics described above the kinetic parameters have to be suitably different to achieve a separation of the kinetics even with depth and overlaying photochromic material. From our experience we deem a difference in $t_{1/2}^{off}$ of ~5 x the sample rate to be enough to distinguish different label populations (in our case of a 10 Hz laser a difference of ~0.4 s). Consequently, higher laser repetition rates will easily allow to distinguish labels with smaller differences effectively expanding the temporal palette for multiplexing.

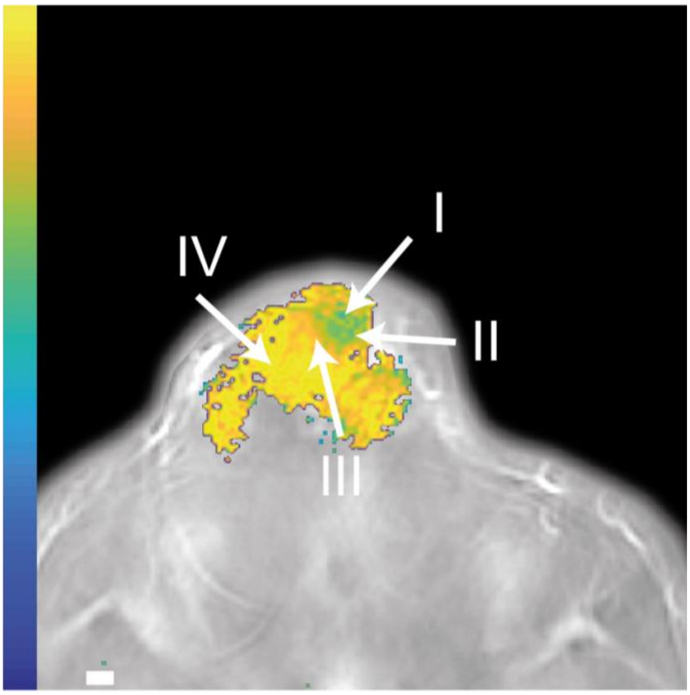


**Suppl. Fig. 8: Concentration dependent change of switching kinetics.** 4T1 tumor at day 14 expressing *Re*BphP-PCM injected with 0.8 x $10^6$ bacteria/µl *E. coli* expressing *Dr*BphP-PCM imaged immediately after injection. Color code represents the kinetics at each pixel. From the center of injection (Arrow I) outward an acceleration of kinetics (Arrow II and III) can be observed suggesting different mixtures between the fast *Re*BphP-PCM (Arrow IV) and the slow *Dr*BphP-PCM (I). Such mixed kinetics suggest that the amount of different labels in a voxel can be inferred from the mixed kinetics. Note that slower kinetics at the border are artefacts due to a low SNR in those areas (due to very low label contend), leading to artificially slow kinetics.

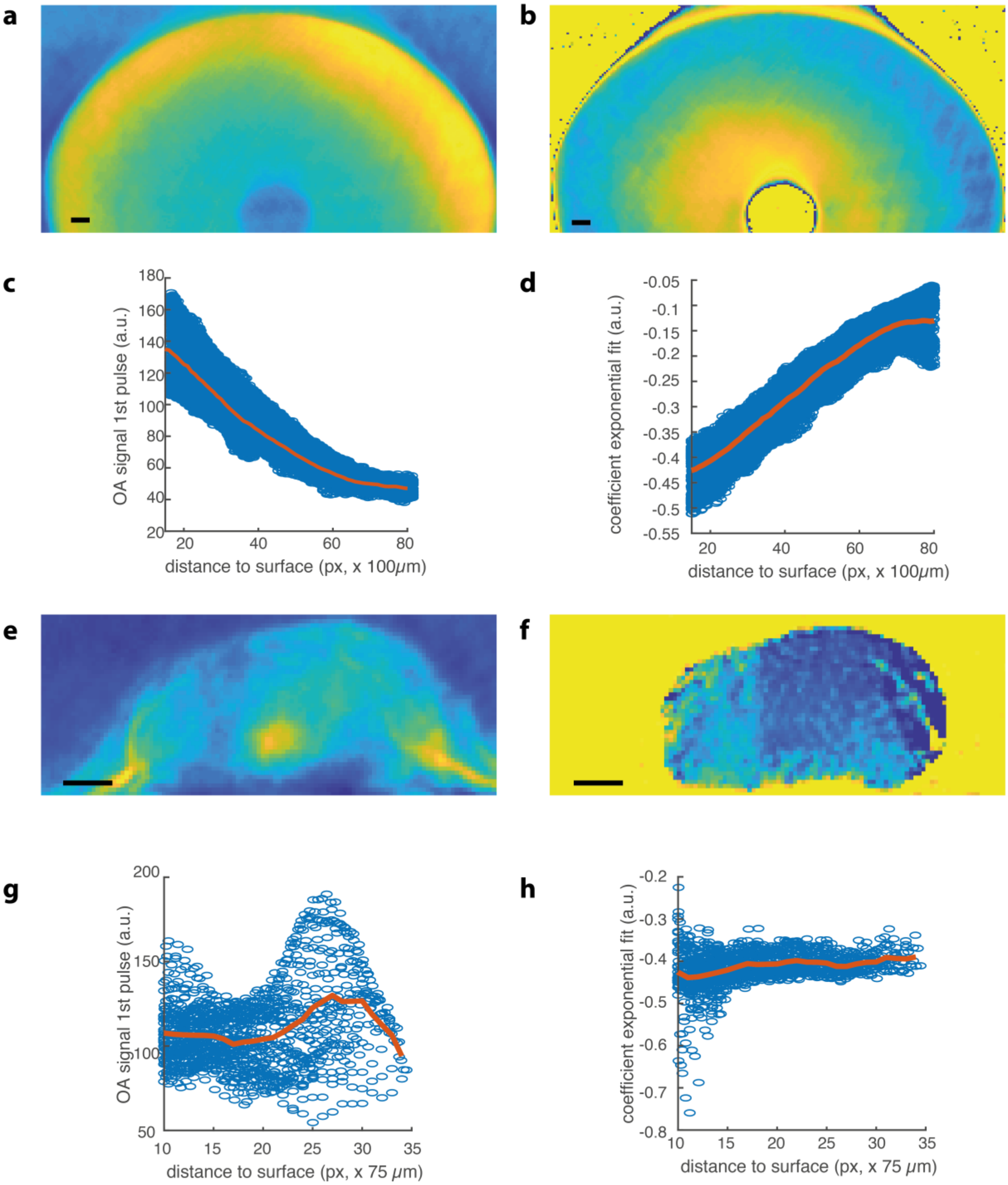


**Suppl. Fig. 9: Dependence of switching kinetics on depth in switchable material** (photochromic protein) for a phantom (**a-d**) and a 4T1 tumor (**e-h**). Shown are the intensity at first pulse (approximately the light fluence, **a and e**) as well as the mean kinetic over all cycles (**b and f**) together with the changes of those with distance to the surface (**c and g** as well as **d and h**).

**Suppl. Material References:**

1. Li, L. *et al.* Small near-infrared photochromic protein for photoacoustic multi-contrast imaging and detection of protein interactions in vivo. *Nat. Commun.* **9**, 2734 (2018).
2. Bellini, D. & Papiz, M. Z. Structure of a Bacteriophytochrome and Light-Stimulated Protomer Swapping with a Gene Repressor. *Structure* **20**, 1436–1446 (2012).
3. Yang, X., Kuk, J. & Moffat, K. Crystal structure of Pseudomonas aeruginosa bacteriophytochrome: photoconversion and signal transduction. *Proc. Natl. …* (2008).
4. Wagner, J. R. *et al.* Mutational analysis of Deinococcus radiodurans bacteriophytochrome reveals key amino acids necessary for the photochromicity and proton exchange cycle of phytochromes. *J. Biol. Chem.* **283**, 12212–26 (2008).
5. Breiman L. Random Forests. *Mach. Learn.* **45**, 5–32 (2001).
6. Burges, C. J. C. A Tutorial on Support Vector Machines for Pattern Recognition. *Data Min. Knowl. Discov.* **2**, 121–167 (1998).
7. Liaw, A. & Wiener, M. Classification and Regression by randomForest. *R News 2* **2**, 18–22 (2002).
8. Dietterich, T. G. An Experimental Comparison of Three Methods for Constructing Ensembles of Decision Trees: Bagging, Boosting, and Randomization. *Mach. Learn.* **40**, 139–157 (2000).
9. Osuna, E., Freund, R. & Girosi, F. *Support Vector Machines: Training and Applications*. (1997).
10. Deán-Ben, X. L. *et al.* Light fluence normalization in turbid tissues via temporally unmixed multispectral optoacoustic tomography. *Opt. Lett.* **40**, 4691–4 (2015).
11. Lihong V. Wang & Wu, H. *Biomedical Optics: Principles and Imaging*. (Wiley-Interscience, 2007).
12. Vetschera, P. *et al.* Characterization of reversibly switchable fluorescent proteins (rsFPs) in optoacoustic imaging. *Anal. Chem.* (2018). doi:10.1021/acs.analchem.8b02599
13. Yao, J. *et al.* Multiscale photoacoustic tomography using reversibly switchable bacterial phytochrome as a near-infrared photochromic probe. *Nat. Methods* **13**, 1–9 (2015).
